# One Method to Sequence Them All? Comparison between Whole-Genome Sequencing (WGS) and Target Enrichment (TE) of museum specimens from the moth families Epicopeiidae and Sematuridae (Lepidoptera)

**DOI:** 10.1101/2023.08.21.553699

**Authors:** Elsa Call, Victoria Twort, Marianne Espeland, Niklas Wahlberg

## Abstract

There are various possibilities for sequencing highly degraded DNA, such as target enrichment (TE), or whole-genome sequencing (WGS). Here we compare TE and WGS methods using old museum specimens of two families of moths in the superfamily Geometroidea: Epicopeiidae and Sematuridae. Until recently, the relationships of these two families were unclear, as few studies had been done. Recently two studies used the TE approach, either on relatively fresh specimens, or on old museum specimens. Here, we aim to increase the sampling of the families Epicopeiidae and Sematuridae from museum specimens using the WGS method. We show that both sequencing methods give comparable results, but, unsurprisingly, WGS recovers more data. By combining TE and WGS data, we confirm that Sematuridae are sister to Pseudobistonidae+Epicopeiidae. Relationships of genera within the families are well supported. With the costs of WGS decreasing, we suggest that using low-coverage whole genome sequencing is becoming an increasingly viable option in the phylogenomics of insects.

## INTRODUCTION

The ability to efficiently sequence an organism’s genome has revolutionised biological sciences. Studying genomes has become a broad research area that covers many aspects; such as systematics, evolution, population genetics, functional genetics, and metagenomics. Due to the wide applicability of genomics approaches, a large panel of methods have been developed. In phylogenomics, three methods are commonly used and yet profoundly different in their approaches. Two of these methods are genome-reduction approaches: transcriptomics and target enrichment, while the third method is whole-genome sequencing (WGS) with various coverage.

Transcriptomics is the study of the expressed RNA in a particular cell or tissue at the point in time of sampling (Wang et al. 2009; Lowe et al. 2017). This technique can be used to advance knowledge of phylogenetic relationships by inferring orthologous coding sequences (Bazinet et al. 2013; Kawahara and Breinholt 2014; Bazinet et al. 2017; Kawahara et al. 2019). However, transcriptomic methods cannot be applied to museum specimens, as they typically require fresh material with RNA still intact (Wang et al. 2009; Lowe et al. 2017).

The second set of techniques, also genome reduction methods, aims to target, amplify and sequence distinct parts of the genome. Two main methods have been developed: target enrichment (TE) (Meyer and Kircher 2010; Lemmon et al. 2012) and ultra-conserved elements (UCE) (Faircloth et al. 2012; McCormack et al. 2012). These methods are particularly useful in phylogenomics, as they focus on specific genomic regions that have been shown to be phylogenetically informative (Kadlec et al. 2017; Toussaint et al. 2018). In particular, TE has been successfully applied to phylogenomics of butterflies and moths (Breinholt et al. 2018; Espeland et al. 2018; Kawahara et al. 2018; Espeland et al. 2019; Homziak et al. 2019; Mayer et al. 2021).

Finally, as the name suggests, whole-genome sequencing (WGS) methods intend to sequence the entire genome of an organism. Today it is relatively simple to sequence a genome using a variety of available platforms, although how much of the genome is actually sequenced depends on the size of the genome and the coverage depth. The challenge lies in the processing of the resulting raw sequencing data, or so-called reads. There are two ways of assembling the data derived from whole genome sequencing (Ng and Kirkness 2010; Lu et al. 2013); the first one, *de novo* assembly, reconstructs longer contiguous sequences from raw reads by comparing sequence reads to each other. When no closely related genome reference sequence exists, this method is the only possible approach (Zerbino and Birney 2008). The second way, reference assembly, is when reads are assembled against an existing reference genome, usually from the same species or one that is very closely related (Church et al. 2011; Lu et al. 2013; Ekblom and Wolf 2014). Despite its advantages, this technique is not as popular in phylogenomics, mainly due to the lack of reference genomes (Savva et al. 2003; Pearson et al. 2004; Croucher et al. 2015; Marcet-Houben and Gabaldon 2015).

Here we compare WGS to TE with regard to museum specimens. Museomics is a field that focuses on the DNA from museum specimens. Natural history museum collections represent a diverse and large biobank of samples, including rare and difficult to collect specimens; and even extinct taxa (Grewe et al. 2021). However, museum specimens present their own challenges, with highly fragmented genomes. Often the only way to obtain sequences from such specimens is by using short read technologies, such as the Illumina platform. The resulting reads tend to range from 50 to 150 bp (Sproul and Maddision 2017; Korlević et al. 2021; Mullin et al. 2022), leading to difficulties with assembly. Recent studies have shown that it is possible to extract phylogenetically useful data from such fragmented genomes using both TE (Espeland et al. 2018; St Laurent et al. 2018; Espeland et al. 2019; Call et al. 2021) and WGS (Cong et al. 2017; Sproul and Maddision 2017; Li et al. 2019; Zhang et al. 2019; Cong et al. 2021; Grewe et al. 2021; Twort et al. 2021; Mullin et al. 2022). However, as museomics is such a young field, there is no consensus on which method to access genomic information is the most suitable. Here we want to provide a comparison of these two major sequencing approaches in a phylogenomics perspective.

We focus our attention on two small families of the superfamily Geometroidea (Lepidoptera): Epicopeiidae and Sematuridae. The relationships, not only within these two clades but also between them and Pseudobistonidae, a close family, were unclear until recently (Minet 2002; Rajaei et al. 2015; Wei and Yen 2017; Wang et al. 2019; Zhang et al. 2020). In a previous study, we conducted a phylogenetic analysis using TE on museum specimens of Epicopeiidae, Sematuridae and Pseudobistonidae (Call et al. 2021). Here, we performed a WGS study on Epicopeiidae and Sematuridae. We extended the species coverage among these families by adding the WGS results to our previous TE analysis. We also add our new data to two other datasets, showcasing the uses of WGS for phylogenetic work. Finally, we compared the usefulness of these two sequencing methods for museum specimens.

## MATERIAL & METHODS

### Taxon sampling

We sampled specimens of Epicopeiidae and Sematuridae from four different museums: Natural History Museum of Denmark (NHMD, Copenhagen, Denmark), Naturalis Biodiversity Center (Naturalis, Leiden, Netherland), Naturhistoriska riksmuseet (NRM, Stockholm, Sweden) and National Museum of Nature and Science (NSMT, Tokyo, Japan). We sampled as many species as possible, with two specimens per species when available. In total, we sampled thirteen species of Epicopeiidae and nine species of Sematuridae (Table 1). Seven specimens did not have dated labels; thus, we are not sure about the age of these specimens. The oldest of the dated specimens is *Homidiana leachi* (Sematuridae) from 1907, provided by NHMD. We extracted DNA from a total of thirty-two museum samples. We were not able to acquire specimens of the genera *Burmeia* (monotypic) and *Mimaporia* (2 species) for the family Epicopeiidae, nor samples of the monotypic *Lonchotura* and *Apoprogones* of the family Sematuridae. An overview of all the specimens in this study is available in Table 1 (see Appendix 1 with pictures of the WGS sequenced specimens at: https://doi.org/10.5281/zenodo.3772305).

**Table 1:** Summary of the specimens sequenced (WGS) for this study. Pictures of all specimens are available at https://doi.org/10.5281/zenodo.3772305.

| Family | Species | ID | Year | Museum | Museum ID |
| --- | --- | --- | --- | --- | --- |
| Epicopeiidae | *Amana angulifera* (Walker, 1855) | PE3 | 1935 | Leiden | RMHN.INS.1099000 |
| Epicopeiidae | *A. angulifera* | TE36 | 1998 | Tokyo | NSMT-I-L-49036 |
| Epicopeiidae | *Chatamla flavescens* (Walker, 1854) | TE37 | 2003 | Tokyo | NSMT-I-L-49037 |
| Epicopeiidae | *Epicopeia hainesii* (Holland, 1889) | PE2 | 2004 | Leiden | RMHN.INS.1098999 |
| Epicopeiidae | *E. mencia* (Moore, 1875) | PE1 | 1928 | Leiden | RMHN.INS.1098998 |
| Epicopeiidae | *E. philenora* (Westwood, 1841) | PE6 | NA | Leiden | RMHN.INS.1099003 |
| Epicopeiidae | *E. polydora* (Westwood, 1841) | EC2 | 1914 | Copenhagen |  |
| Epicopeiidae | *E. polydora* | PE5 | 1920 | Leiden | RMHN.INS.1099002 |
| Epicopeiidae | *Nossa nelcinna* (Moore, 1875) | TE38 | 1971 | Tokyo | NSMT-I-L-49038 |
| Epicopeiidae | *N. nelcinna* | TE39 | 2000 | Tokyo | NSMT-I-L-49039 |
| Epicopeiidae | *N. moorei* (Elwes, 1890) | PE4 | 1936 | Leiden | RMHN.INS.1099001 |
| Epicopeiidae | *N. nagaensis* (Elwes, 1890) | TE40 | 2006 | Tokyo | NSMT-I-L-49040 |
| Epicopeiidae | *Parabraxas davidi* (Oberthür, 1885) | EC1 | 2006 | Copenhagen |  |
| Epicopeiidae | *Psychotrophia endoi* (Inoue, 1992) | TE41 | 2001 | Tokyo | NSMT-I-L-49041 |
| Epicopeiidae | *P. melanargia* (Butler, 1877) | ES1 | NA | Stockholm | NHRS-TOBI000003231 |
| Epicopeiidae | *Schistomitra* *funeralis*<br>(Butler, 1881) | TE42 | 2002 | Tokyo | NSMT-I-L-49042 |
| Sematuridae | *Coronidia erecthea*<br>(Westwood, 1879) | SS1 | NA | Stockholm | NHRS-TOBI000003232 |
| Sematuridae | *C. erecthea* | SC4 | NA | Copenhagen |  |
| Sematuridae | *C. orithea*<br>(Cramer, 1780) | SC2 | 1914 | Copenhagen |  |
| Sematuridae | *Homidiana canace*<br>(Hopffer, 1856) | SC6 | 1917 | Copenhagen |  |
| Sematuridae | *H. canace* | SS2 | NA | Stockholm | NHRS-TOBI000003233 |
| Sematuridae | *H. leachi*<br>(Godart, 1819) | SC1 | 1907 | Copenhagen |  |
| Sematuridae | *H. leachi* | SC5 | 1927 | Copenhagen |  |
| Sematuridae | *Mania diana*<br>(Guénée, 1857) | PS3 | 1926 | Leiden | RMNH.INS.1098997 |
| Sematuridae | *M. diana* | SC3 | 1915 | Copenhagen |  |
| Sematuridae | *M. diana* | SS5 | NA | Stockholm | NHRS-TOBI000003236 |
| Sematuridae | *M. empedocles*<br>(Cramer, 1782) | SS3 | NA | Stockholm | NHRS-TOBI000003235 |
| Sematuridae | *M. empedocles* | SC8 | 1931 | Copenhagen |  |
| Sematuridae | *M. lunus*<br>(Linneaus, 1758) | PS1 | 1966 | Leiden | RMHN.INS.1098995 |
| Sematuridae | *M. lunus* | SC7 | 1930 | Copenhagen |  |
| Sematuridae | *M. lunus* | SS4 | 1990 | Stockholm | NHRS-TOBI000003234 |
| Sematuridae | *M. lunus* | PS2 | 1971 | Leiden | RMHN.INS.1098996 |

Historically, the genus *Mania* has also been referred to as either *Sematura* or *Nothus*. However, these names are considered invalid as the name *Nothus* refers to a genus of Coleoptera (Minet and Scoble 1998); while *Sematura* is younger (published in 1825 by Dalman) than Hübner’s name *Mania* from 1821. Therefore in this study, all specimens that were labelled as *Sematura* will be referred to as *Mania*.

### Sample preparation and DNA extraction

We extracted DNA from the abdomen of the specimens, using the QIAamp DNA Micro kit (Qiagen). We followed the manufacturer’s protocol for isolation of genomic DNA from tissue. We did not crush the samples in order to preserve genitalia. To ensure complete submersion of the larger specimens, it was necessary to either double, triple or quadruple the volume of buffer ATL and proteinase K. All the samples were incubated at 56°C overnight. The volumes of the two subsequent reagents (lysis buffer and ethanol) were adjusted accordingly. The entire volume was passed through a single QIAamp MinElute column per sample. Downstream steps were later carried out in the recommended volumes. We verified the presence of DNA on a 1.2% gel that was run at 105 volts for 2.5 hours.

### Molecular methods and sequencing

Library preparation for WGS followed the protocol of Meyer and Kircher (2010) as modified in Twort et al. (2021). We repaired the DNA with USER (Uracil-Specific Excision Reagent) Enzyme (NEB, USA) for deanimation. The reaction mix consisted of 1x Tango Buffer, dNTPs (100 µM each), 1 mM ATP, 0.5 U/µL T4 PNK and USER Enzyme (NEB, USA). The reaction was incubated for 15 min at 25°C, followed by 5 min at 12°C. We performed a purification of the reaction with the MinElute purification kit (Qiagen), and elution in 22 µL EB buffer. Then the adapter-ligation step followed with a reaction mix containing: 1x T4 DNA Ligase Buffer, 5% PEG-4000, 0.1125 U/µL T4 DNA Ligase, and an adapter mix of P7 and P5 adapters (Meyer and Kircher 2010) 2.5 µM each. Reactions were incubated for 30 min at 22°C. We concluded an additional step of purification with MinElute purification kit (Qiagen). We completed adapter fill-in using the following reaction mix: 1x Thermopol reaction buffer, dNTPs (250 µM each) and 0.3 U/µL Bst Polymerase. The reaction was incubated for 20 min at 37°C, followed by the final heat kill step at 80°C for 20 min. For more details, see Twort et al. (2021).

Indexing of each library was performed with 3 µL of library template and a unique single indexing strategy. The amplification mix consisted of 0.5 µL AccuPrime Pfx DNA Polymerase, 2.5 µL AccuPrime reaction mix, 0.3 µM IS4 primer (Meyer and Kircher 2010) and 0.3 µM of indexing primer. Reactions were amplified under the following conditions: 95°C for 2 min, 18 cycles of 95°C for 15 sec, 60°C for 30 sec and 68°C for 60 sec. The amplification step was carried out in six independent reactions, to avoid amplification bias, and pooled prior to purification. Purification, along with size selection, was performed using Agencourt AMPure XP beads. The resulting libraries were quantified, and quality checked with Quanti-iT™ PicoGreen™ dsDNA assay and with a DNA chip on a BioAnalyzer 2100, respectively. Multiplexed libraries were pooled in equimolar concentrations in a pool of 10 samples and sequenced using the Illumina HiSeq 2500 technology.

### Data assembly and clean up

We checked the quality of the raw reads with FASTQC v0.11.5 (Andrews 2010). Although paired-end reads were generated, the resulting sequencing data was carried forward as single-end. The reason behind this choice is that degraded DNA is likely to randomly ligate together during the adapter ligation step of library preparation, resulting in chimaeras of genomic regions (Willerslev and Cooper 2005). However, the sequencing information contained in the reads is still reliable, and therefore more accurate results are obtained by treating data as single-end (Rowe et al. 2011). We removed reads with ambiguous bases (N’s) from the dataset using Prinseq 0.20.4 (Schmieder and Edwards 2011). We then used Trimmomatic 0.38 (Bolger et al. 2014) to remove low-quality bases from their beginning (LEADING: 3) and end (TRAILING: 3), by removing reads below 30 bp, and by evaluation read quality with a sliding window approach. Quality was measured for sliding windows of four base pairs and had to be higher than 25 on average. Afterwards, we *de novo* assembled the genomes with spAdes v3.13.1 (Bankevich et al. 2012) with k-mer values of 21, 33 and 55.

### Orthologue dataset and alignment

We analysed three sets of gene markers, the markers used in Call et al. (2021), the set used by Zhang et al. (2020), and finally a set of 8 standard gene fragments used in a large number of Sanger sequencing studies of Lepidoptera (Wahlberg and Wheat 2008; Mutanen et al. 2010). The Call et al. (2021) set represented 308 nuclear gene markers with an average length of 2,097 bp (range: 282-14,394). The Zhang et al. (2020) set comprised 89 nuclear gene markers, with very little overlap to the Call et al. set (8 gene fragments). We used the entire coding region of these genes as our reference set and mined them from our fragmented genome assemblies using MESPA v1.3 (Neethiraj et al. 2017) as described in detail by Twort et al. (2021). We screened the identified coding regions to remove those containing internal stop codons. We translated the resulting DNA sequences to amino acid sequences and aligned them to pre-existing reference alignments (Zhang et al. 2020; Call et al. 2021) using MAFFT v7.310 (Katoh and Standley 2013). We used the added fragments and auto options, which keeps existing gaps in the alignments and chooses the most appropriate alignment strategy, respectively. Manual screening of the resulting amino acid alignments was carried out to ensure accuracy, screen for the presence of pseudogenes, reading frame and alignment errors, using Geneious Prime 2020.0.3 (https://www.geneious.com/). The corrected amino acid alignments were converted back into nucleotide alignments using pal2nal v14 (Suyama et al. 2006).

For the standard 8 gene dataset, we used BLAST to retrieve the sequences from the assemblies, as these are all single exons (Wahlberg and Wheat 2008). The assemblies were queried against the gene set using a tBlastn (v2.2.28, (Altschul et al. 1990)) approach (1e-5 cut-off). The output of the BLAST search (coordinates of BLAST hits within each transcriptome) was then used to extract the DNA sequences from each assembly using a set of Python scripts written by Carlos Peña (https://github.com/carlosp420/PyPhyloGenomics) (described in Rota et al. 2022). These were then aligned manually and added to a dataset of all taxa with two or more of the 8 genes available. Also the Zhang et al. taxa (full overlap of genes) and the Call et al. taxa (overlap of only two genes, MDH and wingless) were included.

### Phylogenetic analyses

The final dataset for the Call et al. set consisted of a total of 308 genes, across 72 specimens, with the total size of the dataset being 650,308 aligned nucleotide sites. Respectively for the Zhang et al. set we had 89 genes across 116 specimens, with a total dataset size of 186,757 aligned nucleotide sites. Finally, for the standard gene set we included 136 specimens with up to 8 gene fragments and a total aligned dataset of 6410 bp.

We initially partitioned the data by genes. We performed the model selection using ModelFinder (Kalyaanamoorthy et al. 2017) in IQ-TREE 2.1.2 (Nguyen et al. 2015) with the option –m MFP, which considers all nucleotide models and models of rate heterogeneity (Chernomor et al. 2016) to find the best substitution model for each partition. We then inferred the phylogenetic relationships, using the optimal models, with IQ-TREE 2.1.2 (Nguyen et al. 2015; Chernomor et al. 2016) under the maximum likelihood (ML) criterion. To investigate the robustness of our inferences, we used 1,000 ultrafast bootstrap replicates (Hoang et al. 2018) with the -bnni option to reduce overestimation of support, and 1,000 replicates for SH-aLRT (Guindon et al. 2010).

## RESULTS

### Sequencing and Gene Identification

Library preparation and sequencing was successful for 30 out of the 32 specimens used in this study, and the raw sequence files are available on the NCBI database under the project PRJNA1062115. In the case of the failed samples, one specimen (*Mania diana* PS3) failed during library preparation, while the other (*Homidiana canace* SS2) failed to produce any sequencing data.

Sequencing resulted in an average of 46,265,382 raw paired-end reads per sample. All sequencing data was treated as single-end from this point, due to the high likelihood of chimera formation of fragmented DNA during library preparation (Willerslev and Cooper 2005; Rowe et al. 2011). After read cleaning and quality control, an average of 82,570,123 single-end reads remained per sample. The average sequencing depth is 13X (range 4-24X Table 2). In the case of sample SS3 (*Mania empedocles*) 1,600 raw reads (paired), and a total of 2,571 cleaned reads (single) were obtained (Table 2), therefore, this specimen was excluded from further analysis.

**Table 2:** Summary of the sequencing data per sample.

| ID | Number of raw reads (paired) | Number of cleaned reads (single) | Average depth | Number of scaffolds | Assembly size (bp) | Average scaffold length | N50 |
| --- | --- | --- | --- | --- | --- | --- | --- |
| EC1 | 43,560,424 | 78,923,668 | 15 | 407,237 | 328,812,015 | 807 | 1,692 |
| EC2 | 20,694,421 | 34,043,165 | 4 | 74,44 | 19,712,166 | 265 | 262 |
| ES1 | 36,259,500 | 62,964,513 | 9 | 558,288 | 155,048,062 | 278 | 332 |
| PE1 | 79,647,540 | 142,332,712 | 20 | 497,616 | 281,185,881 | 565 | 894 |
| PE2 | 19,565,912 | 34,438,982 | 6 | 454,461 | 141,199,212 | 311 | 340 |
| PE3 | 15,546,901 | 24,734,523 | 4 | 73,095 | 52,589,789 | 719 | 1,579 |
| PE4 | 21,270,619 | 35,274,799 | 5 | 18,566 | 29,525,395 | 1590 | 9,19 |
| PE5 | 57,804,504 | 101,775,540 | 14 | 456,859 | 265,460,497 | 581 | 752 |
| PE6 | 54,392,683 | 90,375,313 | 21 | 575,571 | 200,907,466 | 349 | 395 |
| PS1 | 60,926,225 | 104,736,878 | 16 | 741,31 | 324,999,839 | 438 | 648 |
| PS2 | 39,707,030 | 67,436,138 | 10 | 610,309 | 215,907,918 | 354 | 402 |
| SC1 | 61,085,459 | 109,315,526 | 15 | 660,594 | 247,843,167 | 375 | 457 |
| SC2 | 54,596,363 | 95,899,455 | 13 | 584,963 | 210,459,262 | 360 | 417 |
| SC3 | 29,617,163 | 52,225,928 | 7 | 210,573 | 54,458,011 | 259 | 270 |
| SC4 | 40,136,635 | 65,301,761 | 8 | 16,883 | 21,820,127 | 1292 | 2,618 |
| SC5 | 55,916,101 | 100,881,708 | 15 | 655,997 | 267,587,339 | 408 | 505 |
| SC6 | 43,770,053 | 78,051,636 | 10 | 493,341 | 159,884,101 | 324 | 356 |
| SC7 | 60,247,023 | 101,518,563 | 13 | 294,086 | 104,819,327 | 356 | 323 |
| SC8 | 88,913,801 | 154,002,434 | 23 | 640,498 | 341,298,631 | 533 | 935 |
| SS1 | 48,950,332 | 85,426,365 | 12 | 595,155 | 220,473,898 | 370 | 418 |
| SS3 | 1,600 | — | — | — | — | — | — |
| SS4 | 60,849,455 | 106,548,432 | 20 | 809,485 | 377,959,964 | 467 | 1,053 |
| SS5 | 53,548,557 | 89,748,410 | 13 | 576,196 | 178,638,636 | 310 | 344 |
| TE36 | 39,067,664 | 64,887,834 | 10 | 688,272 | 255,621,884 | 371 | 439 |
| TE37 | 64,559,940 | 113,796,246 | 24 | 175,741 | 318,398,213 | 1812 | 7,031 |
| TE38 | 44,151,894 | 59,976,931 | 7 | 14,648 | 7,519,867 | 513 | 3,937 |
| TE39 | 64,199,067 | 113,024,824 | 18 | 352,819 | 295,238,493 | 837 | 1,598 |
| TE40 | 38,868,319 | 67,923,489 | 14 | 181,454 | 297,653,559 | 1640 | 2,49 |
| TE41 | 41,034,032 | 71,796,201 | 12 | 807,772 | 298,803,775 | 370 | 612 |
| TE42 | 49,072,254 | 87,171,610 | 19 | 291,448 | 271,378,333 | 931 | 2,976 |

*De novo* assembly resulted in an average assembly size of 205 Mbp (range: 7-377 Mbp). On average, we obtained 431,644 scaffolds (range: 14,648-809,485), with an average N50 of 1,491 (range: 262-9,190) (Table 2, Appendix 2 at: https://doi.org/10.5281/zenodo.3772351). We recovered 308 of the 327 genes used in the Call et al. (2021) analysis using MESPA. Figure 1 compares the distribution of gene lengths recovered by each method for the Call et al. set of genes. The difference in length (bp) recovered between the TE and WGS methods is significant (Df=1, f=468.64, p < 0.001). WGS recovered the longest sequences (7,167 bp Table 3). However, the overall average recovery per sample of the 327 genes used in Call et al. (2021) was better with the TE approach (236 compared to 115 with WGS). We compared what proportion of the reference sequence was recovered for each gene. We recovered, on average, a larger proportion of a given gene with the TE method (24.14%) than with WGS (19.91%).

**Table 2:** Summary of recovered genes length for TE (28 specimens) and WGS (28 specimens) methods.

| Attribute | TE | WGS |
| --- | --- | --- |
| Minimal size (bp) | 6 | 27 |
| 1st Quartile (bp) | 258 | 279 |
| Median (bp) | 297 | 510 |
| Mean (bp) | 336 | 786 |
| 3rd Quartile (bp) | 354 | 1008 |
| Maximal size (bp) | 1752 | 7167 |
| Missing genes/specimen | 2005 | 5394 |
| Average number of genes recovered | 236 | 115 |
| Proportion of the genes recovered on average (%) | 24.14 | 19.91 |
| Min. of the proportion of the recovered genes (%) | 0.23 | 0 |
| Max. of the proportion of the recovered genes (%) | 92.31 | 93.92 |

**Figure 1.**
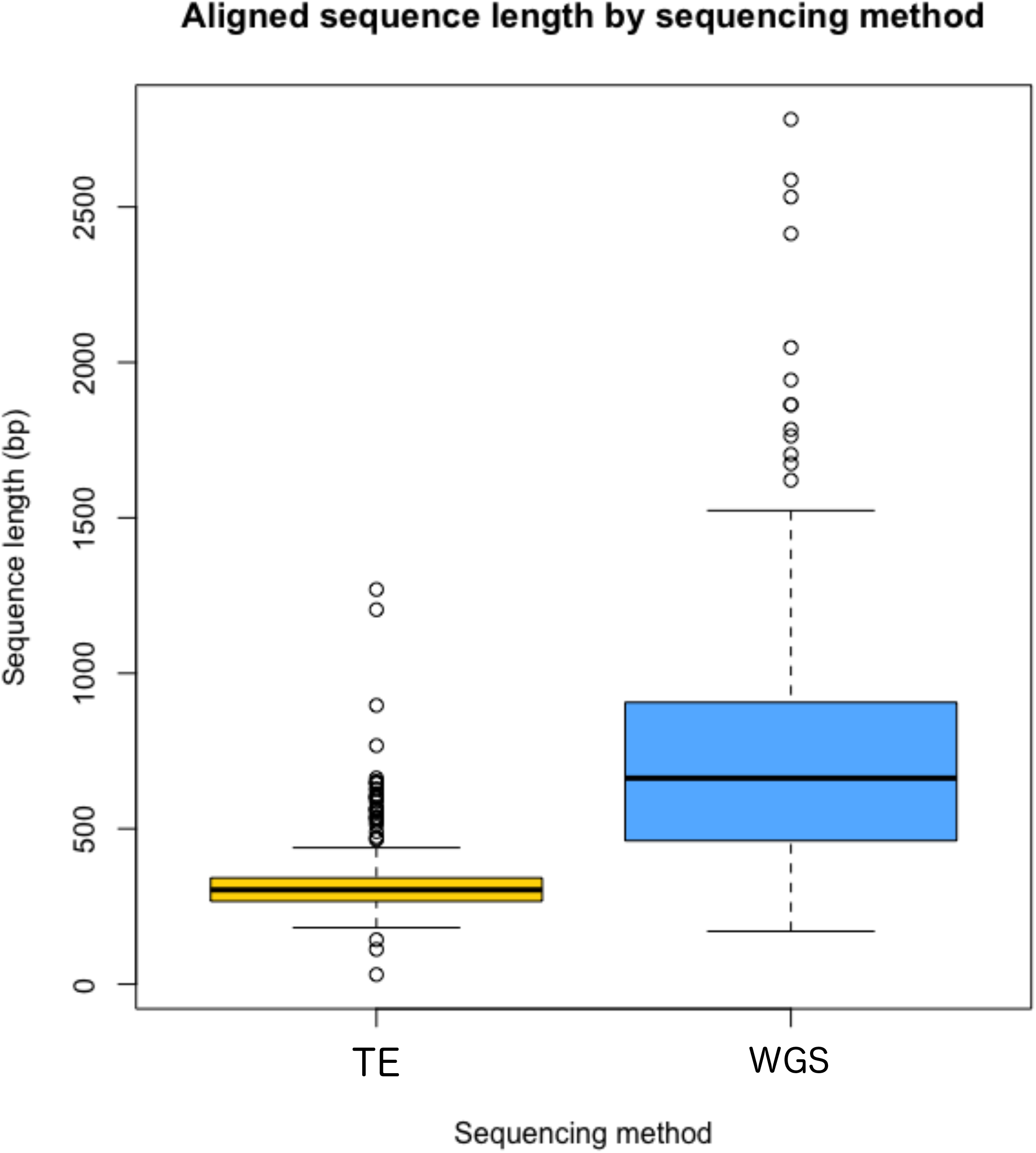
Distribution of the average (bp) recovered per gene by the TE (yellow) and WGS (blue) methods.

The specimens PE3 (*Amana angulifera*), PE4 (*Nossa moorei*), SC4 (*Coronidia erechthea*) and TE38 (*Nossa nelcinna*) were well represented by genes in the final datasets. However, they grouped together in preliminary phylogenetic analyses with much longer branches than the rest of the tree, and blasting the sequences from these specimens on NCBI suggested that they originated from fungi. They were thus excluded from further analyses.

From a number of specimens, we recovered the DNA barcode part of the *COI* sequence and blasted them against the Barcoding Of Life Data (BOLD) system (Ratnasingham and Hebert 2007). Most of them matched to the species they were supposed to be, but SC1 (*Homidiana leachi*) matched *Mania lunus* instead. The identification of SC1 as *Homidiana* is not questioned (see the specimen picture in Appendix 1 available at: https://doi.org/10.5281/zenodo.3772305); thus, there was likely to have been contamination during laboratory work. Furthermore, SC1 was our oldest sampled specimen (with a known collection date), collected in 1907; hence, its DNA must be highly fragmented and in low quantities. Consequently, contamination with any other more recent specimen could have dominated DNA from SC1. Because of this contamination, we did not include this specimen in our analyses.

### Phylogenetic analyses

Our three datasets (Call et al., Zhang et al., and Sanger datasets) had broadly similar results (Figs. 2, 3, S1). We find high support for Epicopeiidae as monophyletic and the sister group of Pseudobistonidae, with Sematuridae being sister to these two with high support. Reassuringly, intraspecific sequences obtained through low-coverage whole genome sequencing are identical or near identical to those obtained using target enrichment and Sanger sequencing for those species with multiple individuals included.

**Figure 2.**
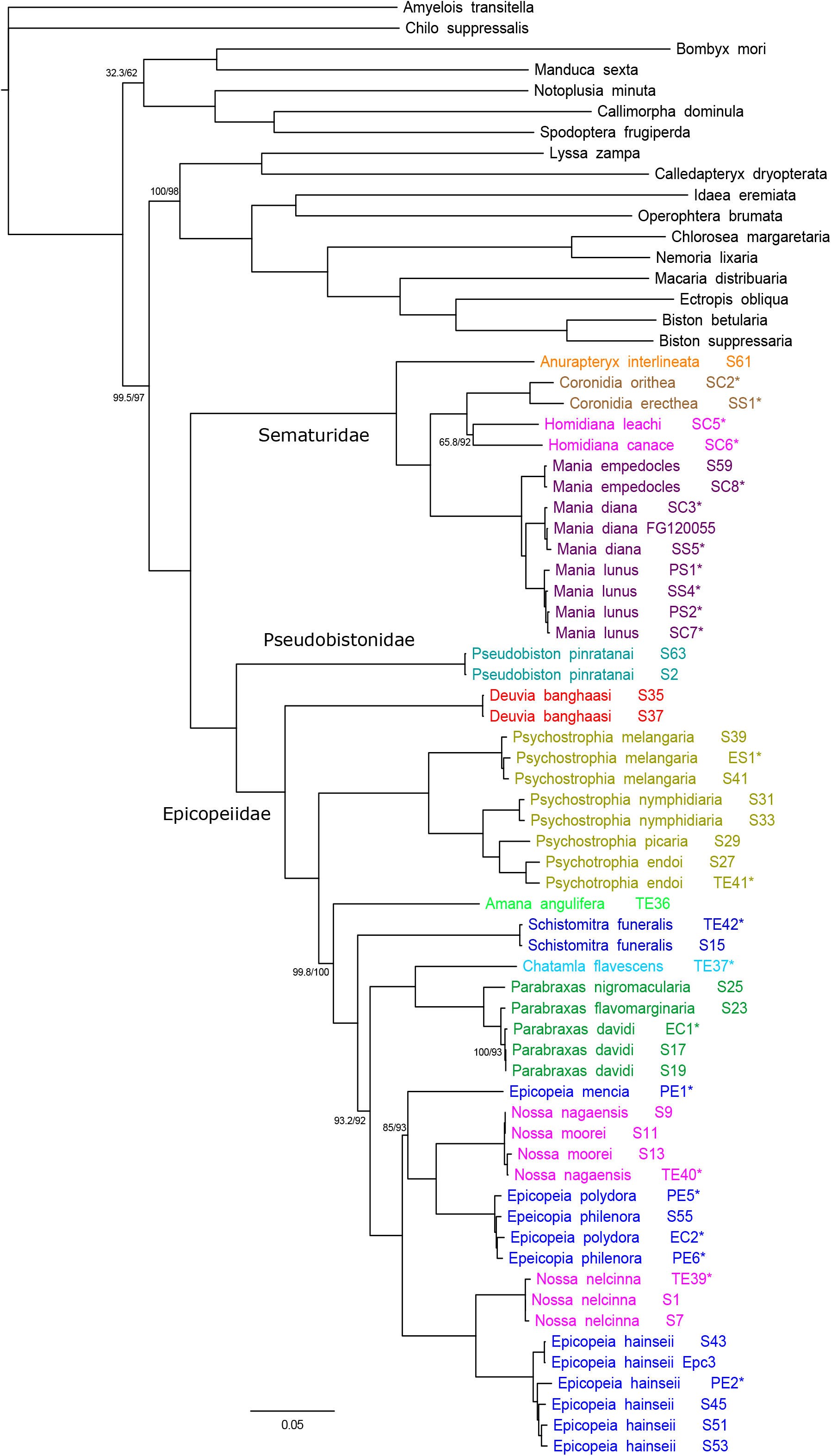
Phylogenetic tree from IQTREE analysis of 45 specimens, based on 308 genes (Call et al. dataset). When displayed, numbers are SH-aLRT support (%) / ultrafast bootstrap support (%). If the support values are equal to 100/100, they are not shown for the nodes. Specimens that have been whole-genome sequenced (WGS, this study) are indicated with an asterisk (*).

**Figure 3.**
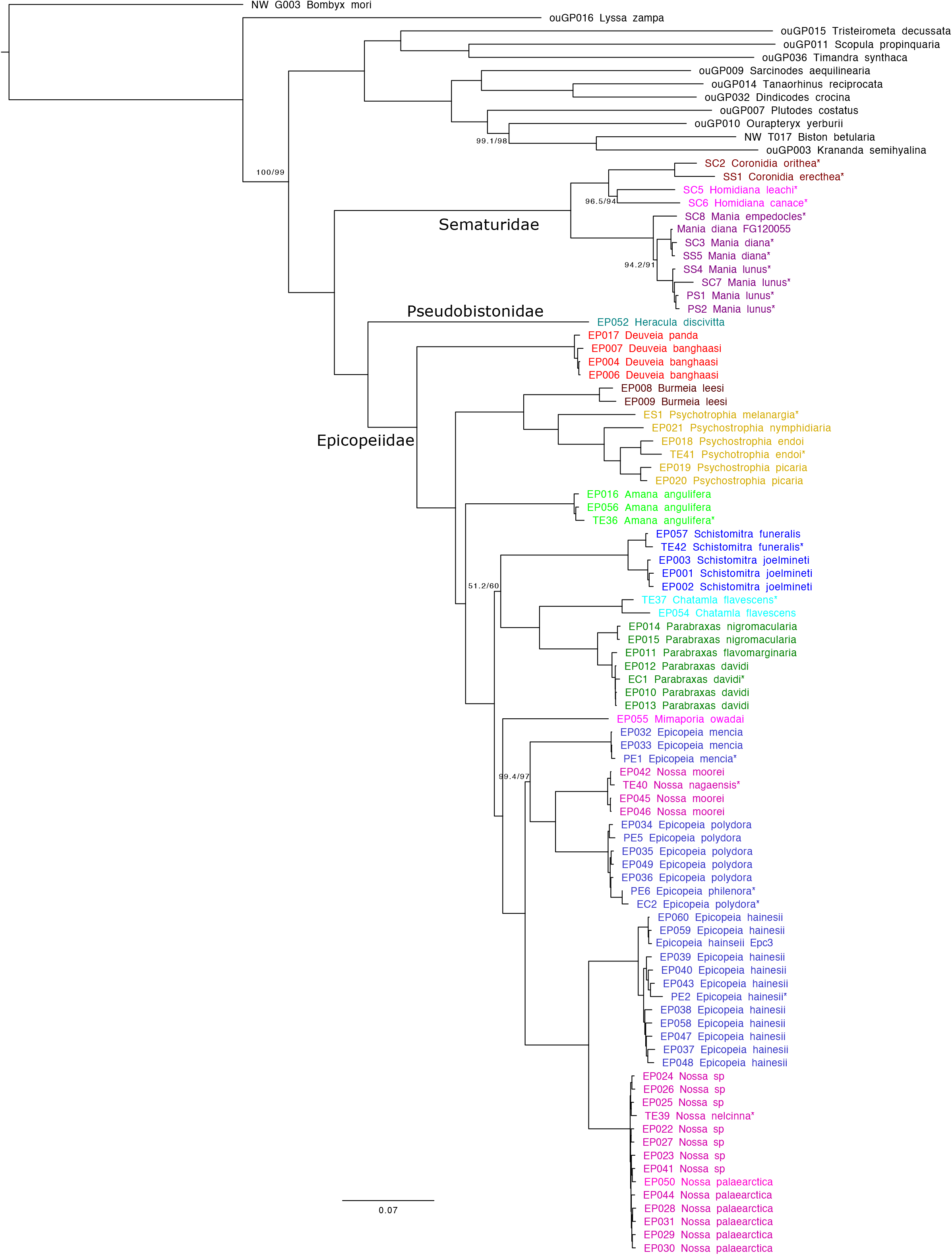
Phylogenetic tree from IQTREE analysis of 96 specimens, based on 89 genes (Zhang et al. dataset). When displayed, numbers are SH-aLRT support (%) / ultrafast bootstrap support (%). If the support values are equal to 100/100, they are not shown for the nodes. Specimens that have been whole-genome sequenced (WGS, this study) are indicated with an asterisk (*).

For Epicopeiidae, the relationships of most genera are mostly congruent with morphological hypotheses (Minet 2002), as well as phylogenetic hypotheses generated from molecular data (Zhang et al. 2020; Call et al. 2021). However, there are some important differences (Figs. 2, 3, S1). The position of *Amana* in the Sanger sequence dataset is sister to *Burmeia+Psychostrophia* (Fig. S1), where as with both genomic approaches, it is sister to a larger clade of of *Schistomitra* to *Epicopeia* (Figs. 2, 3). Also *Schistomitra* has an unstable position, with all three analyses differing from each other. The Call et al. dataset places it sister to *Chatamla+Parabraxas+Nossa+Epicopeia* (Fig. 2), while the Zhang et al. dataset places it sister to *Chatamla+Parabraxas* (Fig. 3), and the Sanger sequence dataset places it sister to *Nossa+Epicopeia* (Fig. S1). The genus *Mimaporia* could not be included in the Call et al. dataset due to the lack of overlapping data, but in the Zhang et al. dataset it is sister to *Nossa+Epicopeia*, while in the Sanger sequence dataset it is within this clade, sister to *Epicopeia hainseii+Nossa palaearctica*.

Within Sematuridae, *Anurapteryx* could only be included in the Call et al. dataset, where it is sister to the rest, *Homidiana* + *Coronidia* are sisters to *Mania. Mania* divides into three clear clades, corresponding to the three sequenced species (Fig. 4). Relationships of taxa common to all datasets are identical.

**Figure 4.**
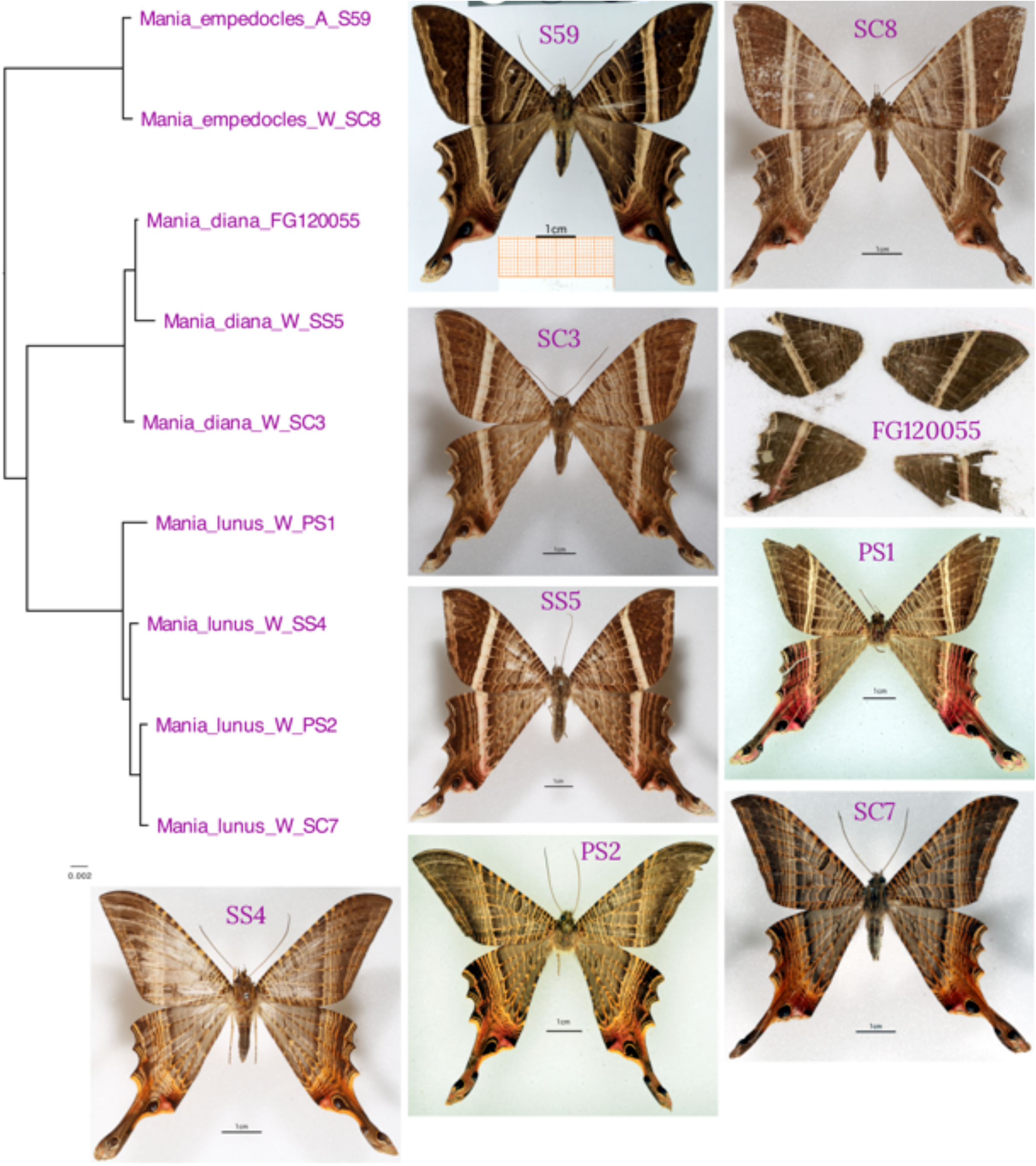
Relationships of species in *Mania*, with pictures of voucher specimens.

Within Pseudobistonidae, *Pseudobiston* and *Heracula* could only be included together in the Sanger sequence dataset due to lack of overlap for the two target enrichment studies. Based on the eight genes, the two are sister species with strong support. Unfortunately neither species had its whole genome sequenced.

## DISCUSSION

### Library preparation, sequencing and gene identification

Of the 32 specimens used for WGS, three samples failed at various stages, which gives a failure rate of 9%. Similarly, our TE study had a failure rate of 6% (Call et al. 2021). In the case of *Mania diana* (PS3), library preparation failed. This failure might be due to several reasons, including for example, low quality or insufficient starting material. This is likely because the sample was collected in 1920, and the DNA is likely to be highly degraded. Other possible reasons are that the DNA extract contained an inhibitor, thereby impeding the successful construction of a library, or that the library may have been lost during the final bead clean. For the samples SS2 (*Homidiana canace*) and SS3 (*Mania empedocles*), the sequencing failed. No sequencing reads were obtained for *H. canace*, while very little data resulted from *M. empedocles*. The potential reasons for this failure are; (i) user error when pooling the samples; (ii) inefficient adapter ligation during library preparation; (iii) ethanol carry over during bead clean-up resulting in library loss or over-drying of the magnetic beads impeding elution. Since we used only a portion of the DNA extract, we would have the possibility to try again to sequence these failed samples.

For the Call et al. dataset, we were able to recover 308 of the 327 TE genes of interest. The length of the recovered genes is significantly longer when using the WGS method than the TE approach for old museum specimens (Figure 1, Table 3). These short fragments (336 bp on average) recovered with TE, tended to be more contiguous, while WGS resulted in longer, but more fragmented gene regions (average length of 786 bp). Similar patterns were observed for the Zhang et al. dataset, although these were not quantified.

In the case of PE3, PE4, SC4 and TE38, these specimens were excluded from downstream analysis. These specimens belong to the following mostly unrelated species, *Amana angulifera, Nossa moorei, Coronidia erechthea* and *Nossa nelcinna*, respectively. After manual screening of the alignments, the gene fragments identified with MESPA generally appeared to be similar to each other, but highly dissimilar to the genes from all other specimens. Comparing these sequences to the NCBI database using BLAST suggested that they were from fungi that had invaded the specimens after being collected.

Regarding *Homidiana leachi* (SC1), whose *COI* sequence matched with *Mania lunus*. This is likely a contamination error that may have occurred during DNA extraction. As previously mentioned, this specimen was the oldest known of our sample. Therefore, cross-contamination with more recent material can dominate the extract.

### Phylogenetic relationships across taxa

*Pseudobiston pinratanai* was recently placed into its own family Pseudobistonidae by Rajaei et al. (2015) who suggested that it was the sister group of Epicopeiidae. Subsequently, another species, *Heracula discivitta*, was added to the family (Wang et al. 2019), and the supposed position corroborated. However, both of these studies were based on eight genes, had few representative species for Epicopeiidae and Sematuridae (three and two, respectively), and had low support for the nodes of those relationships. In their study based on 94 gene fragments, Zhang et al. (2020) found *Heracula* to be sister to Epicopeiidae with strong support. Similarly, Call et al. (2021) showed that *Pseudobiston* was the sister group of Epicopeiidae, and Sematuridae as the sister group to the other two, with strong support. Our current analysis corroborates the monophyly of Pseudobistonidae and its position as sister to Epicopeiidae, reinforcing the case for the new family.

### Phylogenetic relationships among Sematuridae

We show for Sematuridae that *Anurapteryx* is the sister group to the rest of the family. We also show that *Homidiana* and *Coronidia* form a clade that is sister to *Mania*. Within *Mania*, we find *M. empedocles* is sister to *M. diana* + *M. lunus* (Fig. 4). We are only missing *M. aegisthus*, which is endemic to Jamaica, from this genus. Initially, we found that several *Mania* specimens in all the sampled collections were misidentified, and most of these were either *M. empedocles* or *M. diana* lumped under *M. lunus*. Indeed, Sematuridae is not a well studied family, and there is no recent literature to identify specimens. In order to verify identifications, we compared our sampled specimens to pictures of the type specimens (either a photograph in the case of *M. lunus* or original paintings for the other two species). We note that the published transcriptome of a *Mania* species (FG120055, SRA accession SRR1299318) (Kawahara and Breinholt 2014; Kawahara et al. 2019) belongs to the species *M. diana*, not *M. lunus* as reported (see Fig. 4).

### Phylogenetic relationships within Epicopeiidae

Regarding the family Epicopeiidae, our results are largely congruent with each other as well as with the pioneering hypothesis of Minet (2002). Our results continue to be strongly incongruent with Wei and Yen’s (2017) poorly supported results, which is not surprising given that their results were based on only 3 gene fragments. As in the two recent molecular studies (Zhang et al. 2020; Call et al. 2021), we showed *Deuveia* to be sister to the rest of Epicopeiidae. The monophyletic *Psychostrophia* is clearly the sister to *Burmeia*, although this taxon was only available for the Zhang et al data. *Amana* was not included in Call et al. (2021), and when included here it comes out in the same place as the Zhang et al. data suggests, i.e. sister to a large clade comprising *Schistomitra* to *Epicopeia* (Figs. 2 and 3). The Sanger dataset does not show the same pattern, and instead places *Amana* as sister to *Burmeia+Psychostrophia*, albeit with lower support. None of the analyses support Minet’s (2002) hypothesis that *Amana* is sister *Chatamla+Parabraxas*. This can be seen as an illustration of how more data can resolve the relationships of some taxa with more confidence. *Parabraxas* and *Chatamla* are sister taxa, as Minet suggested in 2002, but the position of *Schistomitra* is not consistent in any of the analyses, with only the Sanger dataset being consistent with Minet’s (2002) hypothesis (Fig. S1). Otherwise *Schistomitra* is either placed sister to *Chatamla+Parabraxas* (Zhang et al. dataset, Fig. 3), or as sister to a larger clade of *Chatamla* to *Epicopeia* (Call et al. dataset, Fig. 2). In all cases support is very low for the position of *Schistomitra. Schistomitra* is an example of more data not always allowing confident placement of some taxa, in contrast to the case of *Amana*.

### *Epicopeia* and *Nossa*: a mysterious clade

*Epicopeia* and *Nossa* are polyphyletic with regard to each other (Figs. 2, 3), a result consistent with Wei and Yen (2017), Zhang et al. (2020), and Call et al. (2021). This is surprising as *Nossa* species are superficially similar to each other (mimicking pierid butterflies), as are *Epicopeia* species (mimicking papilionid butterflies), with the biggest difference being the hindwing tails of the latter genus. However, Zhang et al. (2020) discuss some morphological features that might support the molecular results. These characters were, however, not deemed strong enough by the authors to synonymize the genera. The newly described genus *Mimaporia* is found to be sister to *Epicopeia+Nossa* with the Zhang et al. data, as was reported by Zhang et al. (2020), but the Sanger dataset places it within the *Epicopeia+Nossa* clade, sister to *E. hainseii+N. nelcinna* (Fig. 5). This is likely the result of a much smaller dataset for the Sanger sequences.

The species complexes within the *Epicopeia+Nossa* clade are in need of revision. Our results suggest that *E. polydora* (a widespread species in SE Asia) and *E. philenora* (a rare species found only in NE India) are genetically very similar and may represent the same species. Likewise, *Nossa nagaensis* and *N. moorei* (both found in SE Asia) are genetically identical based on the markers we used, suggesting that they are one species. Indeed *N. nagaensis* has recently been synonymised with *N. moorei* based on indistinguishable male genitalia (Kishida 2020). Also the *Nossa nelcinna* complex is in need of revision, names that are involved are *N. palaearctica, N. chinensis*, and *N. leechii*.

### TE vs WGS: the more, the merrier

As mentioned before, the length of the aligned sequences is more than twice the size for the WGS method than the TE approach with regard to museum specimens. With the TE method, we are fairly certain to obtain the probe region of the desired genes and often part of the flanking regions. However, the regions we get from this approach are short (336 bp on average). With WGS, in principle, we should be able to recover the entire gene, and indeed with museum specimens, we do get broader parts of the genes (786 bp on average). However, not all genes were found in the WGS data. Indeed, it is less probable to get the targeted genes in old specimens with highly fragmented DNA, but if present, they are usually longer. Increasing sequencing coverage might also increase the number of genes sequenced, but this also increases the sequencing costs per sample.

On average, we recovered 24% of the reference gene length for TE, while for WGS we recovered 20% (Table 2). The recovery length is similar for both approaches, despite the fact that the gene set being investigated was biased towards the TE data, as we have only investigated the genes that had an efficient TE capture rate. Further analysis, identifying all 2,000 orthologues used in the TE approach from the WGS data would provide a better comparison of the recovery rates and gene length recovery. Interestingly, the recovery rate is similar for both approaches. WGS has the added advantage of being able to go back to the data to mine for new genes of interest that are not included in the TE probes.

Target enrichment approaches have now been used in a number of studies on Lepidoptera (among other taxa), showcasing their utility at various levels of phylogenetic depth (Kadlec et al. 2017; Espeland et al. 2018; Kawahara et al. 2018; St Laurent et al. 2018; Toussaint et al. 2018; Espeland et al. 2019; Homziak et al. 2019; Mayer et al. 2021). They have been used successfully for both fresh material as well as museum specimens. However, as stated above, this approach requires designing specific probes and focuses only on specific parts of the genome. Moreover, in order for probes to capture loci across a large phylogenetic range; they are usually designed against highly conserved regions of the genome. Although with WGS, we will be able to find more genetic information that is informative at very shallow phylogenetic level, this can also be achieved by using flanking regions of the targeted genes (TE). Indeed, TE generally sequences the probe regions and at least a few hundred bps at each end of these, depending on the insert size. Thus, the fact that probes are conserved regions is not a problem since more variable data where this is needed. However, for museum specimens, we showed here and in Call et al. (2021), that the flanking regions of the probes usually do not pass the cleaning process. This is, however, not always the case, and the flanking regions can be useful for museum specimens as shown by Mayer et al. (2021). Therefore, in the case of *Epicopeia* and *Nossa*, for example, obtaining additional genetic information, with TE will require more intensive preparation (e.g. adding more loci to the probe kit). Hence, it can be a great asset to have access to more data (WGS) than targeted genes (TE).

Another advantage of WGS is highlighted by our study. Previously two independent TE based studies had been done on Epicopeiidae (Zhang et al. 2020; Call et al. 2021), but data from these two studies could not be combined as there was very little overlap in the gene regions that were targeted. Our WGS data could be easily added to both datasets, and indeed to any other phylogenomic datasets that are available.

Another asset to consider is that museum specimens already have degraded DNA (Staats et al. 2013; Korlević et al. 2021), therefore, genome reduction methods signify another level of data loss. In short, both approaches have their advantages and disadvantages. However, given the major advances in both sequencing and bioinformatic techniques, we believe a low-coverage whole genome approach will be seen as the more advantageous route in the near future for museum specimens. At the very least, WGS allows for many more questions to be addressed with the same data.

## CONCLUSIONS

We successfully generated WGS data from museum specimens of two small and rarely collected families of Lepidoptera (Epicopeiidae and Sematuridae). We were able to assemble genome data of approximately 205 Mbp in size. We were able to recover 308 of the 327 genes used in a previous TE analysis (Call et al. 2021). We showed a significant difference in the length (bp) of the recovered genes between the TE and WGS methods. Hence, we demonstrate that the WGS approach can recover longer, but more fragmented, sequences from old museum specimens.

Finally, our phylogenetic analysis corroborated the monophyly of Epicopeiidae, as the sister group of Pseudobistonidae, with Sematuridae as the sister clade of these two. Our study shows strong support for *Anurapteryx* being the sister to the rest of our sampled Sematuridae, as well as *Homidiana* and *Coronidia* together as a sister group of *Mania*. The three sampled *Mania* species form genetically separated clades that correspond to their species. Within Epicopeiidae, despite the incongruence regarding the position of *Schistomitra*, our results are congruent with Minet’s (2002) hypothesis, with stronger support than our previous study (Call et al. 2021). We corroborated the polyphyly of both *Epicopeia* and *Nossa* concerning each other. Further studies on these genera are needed to understand the patterns we observed.

Genome reduction methods are a valuable part of our current toolkit to help investigate and understand the phylogenetic relationships of non-model organisms. However, one should keep in mind that, depending on the situation, such sequencing techniques are not always the most suitable or best approach, especially when considering museum specimens. We believe WGS is the way to go for fragmented genomic data, like museum specimens. Furthermore, regarding rare specimens, it is more desirable to obtain as much data as possible, including data that might be useful in the future. For such valuable specimens, increasing the sequencing coverage, and not thinking about cost-efficiency, could further increase the amount of generated data.

## ACKNOWLEDGEMENTS

We are grateful to the curators and museum assistants of the following natural history museums for giving us access to their specimens for this project: Natural History Museum of Denmark (NHMD, Copenhagen, Denmark), Naturalis Biodiversity Center (Naturalis, Leiden, Netherlands), Naturhistoriska riksmuseet (NRM, Stockholm, Sweden) and National Museum of Nature and Science (NSMT, Tokyo, Japan). We are grateful to Rob de Vos, who photographed the specimens from Leiden for us. We thank Akito Kawahara for sending a picture of the voucher specimen of *Mania* used in their studies. The authors acknowledge support from the National Genomics Infrastructure in Stockholm funded by Science for Life Laboratory, the Knut and Alice Wallenberg Foundation and the Swedish Research Council, and NAISS for assistance with massively parallel sequencing and access to the UPPMAX computational infrastructure.

## Funding

This project has received funding from the European Union’s Horizon 2020 research and innovation program under the Marie Sklodowska-Curie grant agreement No. 642241.

## Conflict of interest disclosure

The authors declare no conflicts of interest.

## Data, script, code, and supplementary information availability

Pictures of the WGS sequenced specimens are available at: https://doi.org/10.5281/zenodo.3772305. Raw sequence data are available on the NCBI database under the project PRJNA1062115. Analysed datasets are available at https://doi.org/10.5281/zenodo.8275406. Custom scripts for extracting DNA sequences from BLAST output are available at https://github.com/carlosp420/PyPhyloGenomics.

## Figure legends

**Figure S1**. Phylogenetic tree from IQTREE analysis of 134 specimens, based on 8 genes (Sanger sequence dataset). When displayed, numbers are SH-aLRT support (%) / ultrafast bootstrap support (%). If the support values are equal to 100/100, they are not shown for the nodes. Specimens that have been whole-genome sequenced (WGS, this study) are indicated with an asterisk (*).

## Notes

https://doi.org/10.5281/zenodo.3772305

### Competing Interest Statement

The authors have declared no competing interest.

### Summary of Updates

The manuscript has been revised to include information about the availability of the raw data as well as the analysed datasets. Also an explicit statement of received funding has been included.

## References

Altschul S.F., Gish W., Miller W., Myers E.W., Lipman D.J. 1990. Basic local alignment search tool. Journal of Molecular Biology, 215:403–410. doi.

Andrews S. 2010. FastQC: a quality control tool for high throughput sequence data. http://www.bioinformatics.babraham.ac.uk/projects/fastqc.

Bankevich A., Nurk S., Antipov D., Gurevich A.A., Dvorkin M., Kulikov A.S., Lesin V.M., Nikolenko S.I., Pham S., Prjibelski A.D., et al. 2012. SPAdes: A new genome assembly algorithm and its applications to single-cell sequencing. J. Comp. Biol., 19:455–477. doi: 10.1089/cmb.2012.0021.

Bazinet A.L., Cummings M.P., Mitter K.T., Mitter C.W. 2013. Can RNA-Seq resolve the rapid radiation of advanced moths and butterflies (Hexapoda: Lepidoptera: Apoditrysia)? An exploratory study. PLoS One, 8:e82615. doi: 10.1371/journal.pone.0082615.

Bazinet A.L., Mitter K.T., Davis D.R., van Nieukerken E.J., Cummings M.P., Mitter C. 2017. Phylotranscriptomics resolves ancient divergences in the Lepidoptera. Syst. Entom., 42:305–316. doi: 10.1111/syen.12217.

Bolger A.M., Lohse M., Usadel B. 2014. Trimmomatic: A flexible trimmer for Illumina sequence data. Bioinformatics, 30:2114–2120. doi: 10.1093/bioinformatics/btu170.

Breinholt J.W., Earl C., Lemmon A.R., Lemmon E.M., Xiao L., Kawahara A.Y. 2018. Resolving relationships among the megadiverse butterflies and moths with a novel pipeline for anchored phylogenomics. Syst. Biol., 67:78–93. doi: 10.1093/sysbio/syx048.

Call E., Mayer C., Dietz L., Twort V.G., Wahlberg N., Espeland M. 2021. Museomics: phylogenomics of the moth family Epicopeiidae (Lepidoptera) using target enrichment. Insect Systematics & Diversity, 5:6. doi: 10.1093/isd/ixaa021.

Chernomor O., von Haeseler A., Minh B.Q. 2016. Terrace aware data structure for phylogenomic inference from supermatrices. Syst. Biol., 65:997–1008. doi: 10.1093/sysbio/syw037.

Church D.M., Schneider V.A., Graves T., Auger K., Cunningham F., Bouk N., Chen H.C., Agarwala R., McLaren W.M., Ritchie G.R.S., et al. 2011. Modernizing reference genome assemblies. Plos Biology, 9:e1001091. doi: 10.1371/journal.pbio.1001091.

Cong Q., Shen J., Borek D., Robbins R.K., Opler P.A., Otwinowski Z., Grishin N.V. 2017. When COI barcodes deceive: Complete genomes reveal introgression in hairstreaks. Proc. R. Soc. B, 284:20161735. doi.

Cong Q., Shen J., Zhang J., Li W., Kinch L.N., Calhoun J.V., Warren A.D., Grishin N.V. 2021. Genomics reveals the origins of ancient specimens. Mol. Biol. Evol., 38:2166–2176. doi: 10.1093/molbev/msab013.

Croucher N.J., Page A.J., Connor T.R., Delaney A.J., Keane J.A., Bentley S.D., Parkhill J., Harris S.R. 2015. Rapid phylogenetic analysis of large samples of recombinant bacterial whole genome sequences using Gubbins. Nucleic Acids Research, 43:e15. doi: 10.1093/nar/gku1196.

Ekblom R., Wolf J.B.W. 2014. A field guide to whole-genome sequencing, assembly and annotation. Evolutionary Applications, 7:1026–1042. doi.

Espeland M., Breinholt J., Willmott K.R., Warren A.D., Vila R., Toussaint E.F.A., Maunsell S.C., Aduse-Poku K., Talavera G., Eastwood R., et al. 2018. A comprehensive and dated phylogenomic analysis of butterflies. Curr. Biol., 28:P770–P778. doi: 10.1016/j.cub.2018.01.061.

Espeland M., Breinholt J.W., Barbosa E.P., Casagrande M.M., Huertas B., Lamas G., Marin M.A., Mielke O.H.H., Miller J.Y., Nakahara S., et al. 2019. Four hundred shades of brown: Higher level phylogeny of the problematic Euptychiina (Lepidoptera, Nymphalidae, Satyrinae) based on hybrid enrichment data. Molecular Phylogenetics and Evolution, 131:116–124. doi: 10.1016/j.ympev.2018.10.039.

Faircloth B.C., McCormack J.E., Crawford N.G., Harvey M.G., Brumfield R.T., Glenn T.C. 2012. Ultraconserved ezlements anchor thousands of genetic markers spanning multiple evolutionary timescales. Syst. Biol., 61:717–726. doi: 10.1093/sysbio/sys004.

Grewe F., Kronforst M.R., Pierce N.E., Moreau C.S. 2021. Museum genomics reveals the Xerces blue butterfly (Glaucopsyche xerces) was a distinct species driven to extinction. Biology Letters, 17:20210123. doi: 10.1098/rsbl.2021.0123.

Guindon S., Dufayard J.-F., Lefort V., Anisimova M., Hordijk W., Gascuel O. 2010. New algorithms and methods to estimate Maximum-Likelihood phylogenies: Assessing the performance of PhyML 3.0. Syst. Biol., 59:307–321. doi: 10.1093/sysbio/syq010.

Hoang D.T., Chernomor O., von Haeseler A., Minh B.Q., Vinh L.S. 2018. UFBoot2: Improving the ultrafast bootstrap approximation. Mol. Biol. Evol., 35:518–522. doi: 10.1093/molbev/msx281.

Homziak N.T., Breinholt J.W., Branham M.A., Storer C.G., Kawahara A.Y. 2019. Anchored hybrid enrichment phylogenomics resolves the backbone of erebine moths. Molecular Phylogenetics and Evolution, 131:99–105. doi.

Kadlec M., Bellstedt D.U., Le Maitre N.C., Pirie M.D. 2017. Targeted NGS for species level phylogenomics: “made to measure” or “one size fits all”? PeerJ, 5:e3569. doi: 10.7717/peerj.3569.

Kalyaanamoorthy S., Minh B.Q., Wong T.K.F., von Haeseler A., Jermiin L.S. 2017. ModelFinder: Fast model selection for accurate phylogenetic estimates. Nature Methods, 14:587–589. doi: 10.1038/nmeth.4285.

Katoh K., Standley D.M. 2013. MAFFT Multiple sequence alignment software version 7: improvements in performance and usability. Mol. Biol. Evol., 30:772–780. doi: 10.1093/molbev/mst010.

Kawahara A.Y., Breinholt J.W. 2014. Phylogenomics provides strong evidence for relationships of butterflies and moths. Proc. R. Soc. B, 281:20140970. doi: 10.1098/rspb.2014.0970.

Kawahara A.Y., Breinholt J.W., Espeland M., Storer C., Plotkin D., Dexter K.M., Toussaint E.F.A., St Laurent R.A., Brehm G., Vargas S., et al. 2018. Phylogenetics of moth-like butterflies (Papilionoidea: Hedylidae) based on a new 13-locus target capture probe set. Molecular Phylogenetics and Evolution, 127:600–605. doi: 10.1016/j.ympev.2018.06.002.

Kawahara A.Y., Plotkin D., Espeland M., Meusemann K., Toussaint E.F.A., Donath A., Gimnich F., Frandsen P.B., Zwick A., dos Reis M., et al. 2019. Phylogenomics reveals the evolutionary timing and pattern of butterflies and moths. PNAS, 116:22657–22663. doi: 10.1073/pnas.1907847116.

Kishida Y. 2020. Moths of Laos. Part 1. Tokyo, Japan, Japan Heterocerists’ Society.

Korlevic P., McAlister E., Mayho M., Makunin A., Flicek P., Lawniczak M.K.N. 2021. A minimally morphologically destructive approach for DNA retrieval and whole-genome Shotgun sequencing of pinned historic dipteran vector species. Genom. Biol. Evol., 13:evab226. doi: 10.1093/gbe/evab226.

Lemmon A.R., Emme S.A., Lemmon E.M. 2012. Anchored hybrid enrichment for massively high-throughput phylogenomics. Syst. Biol., 61:727–744. doi.

Li W., Cong Q., Shen J., Zhang J., Hallwachs W., Janzen D.H., Grishin N.V. 2019. Genomes of skipper butterflies reveal extensive convergence of wing patterns. PNAS, 116:6232–6237. doi: 10.1073/pnas.1821304116.

Lowe R., Shirley N., Bleackley M., Dolan S., Shafee T. 2017. Transcriptomics technologies. Plos Computational Biology, 13:e1005457. doi: 10.1371/journal.pcbi.1005457.

Lu B.X., Zeng Z.B., Shi T.L. 2013. Comparative study of de novo assembly and genome-guided assembly strategies for transcriptome reconstruction based on RNA-Seq. Science China-Life Sciences, 56:143–155. doi: 10.1007/s11427-013-4442-z.

Marcet-Houben M., Gabaldon T. 2015. Beyond the Whole-Genome Duplication: Phylogenetic Evidence for an Ancient Interspecies Hybridization in the Baker’s Yeast Lineage. Plos Biology, 13:e1002220. doi: 10.1371/journal.pbio.1002220.

Mayer C., Dietz L., Call E., Kukowka S., Martin S., Espeland M. 2021. Adding leaves to the Lepidoptera tree: capturing hundreds of nuclear genes from old museum specimens. Syst. Entom., in press. doi: 10.1111/syen.12481.

McCormack J.E., Faircloth B.C., Crawford N.G., Gowaty P.A., Brumfield R.T., Glenn T.C. 2012. Ultraconserved elements are novel phylogenomic markers that resolve placental mammal phylogeny when combined with species-tree analysis. Genome Res., 22:746–754. doi: 10.1101/gr.125864.111.

Meyer M., Kircher M. 2010. Illumina sequencing library preparation for highly multiplexed target capture and sequencing. Cold Spring Harbor Protocols, 2010:pdb.prot5448. doi: 10.1101/pdb.prot5448.

Minet J. 2002. The Epicopeiidae: phylogeny and a redefinition, with the description of new taxa (Lepidoptera: Drepanoidea). Annales de la Société entomologique de France, 38:463–487. doi.

Minet J., Scoble M. 1998. The Drepanoid / Geometroid assemblage. In: Kristensen NP editor. Lepidoptera, Butterflies and Moths. Vol. 1. Evolution, Systematics and Biogeography. New York, NY, USA, Walter de Gruyter Inc, p. 301–320.

Mullin V.E., Stephen W., Arce N.A., Nash W., Raine C., Notton D.G., Whiffin A., Blagderov V., Gharbi K., Hogan J., et al. 2022. First large-scale quantification study of DNA preservation in insects from natural history collections using genome-wide sequencing. Meth. Ecol. Evol., in press. doi: 10.1111/2041-210X.13945.

Mutanen M., Wahlberg N., Kaila L. 2010. Comprehensive gene and taxon coverage elucidates radiation patterns in moths and butterflies. Proc. R. Soc. B, 277:2839–2848. doi: 10.1098/rspb.2010.0392.

Neethiraj R., Hornett E.A., Hill J.A., Wheat C.W. 2017. Investigating the genomic basis of discrete phenotypes using a Pool-Seq-only approach: New insights into the genetics underlying colour variation in diverse taxa. Molecular Ecology, 26:4990–5002. doi: 10.1111/mec.14205.

Ng P.C., Kirkness E.F. 2010. Whole genome sequencing. In: M. B, G. B editors. Genetic Variation. Methods in Molecular Biology (Methods and Protocols). Totowa, NJ, USA, Humana Press, p. 215–226.

Nguyen L.-T., Schmidt H.A., von Haeseler A., Minh B.Q. 2015. IQ-TREE: A fast and effective stochastic algorithm for estimating maximum likelihood phylogenies. Mol. Biol. Evol., 32:268–274. doi: 10.1093/molbev/msu300.

Pearson T., Busch J.D., Ravel J., Read T.D., Rhoton S.D., U’Ren J.M., Simonson T.S., Kachur S.M., Leadem R.R., Cardon M.L., et al. 2004. Phylogenetic discovery bias in Bacillus anthracis using single-nucleotide polymorphisms from whole-genome sequencing. PNAS, 101:13536–13541. doi: 10.1073/pnas.0403844101.

Rajaei H.S., Greve C., Letsch H., Stüning D., Wahlberg N., Minet J., Misof B. 2015. Advances in Geometroidea phylogeny, with characterization of a new family based on Pseudobiston pinratanai (Lepidoptera, Glossata). Zool. Scripta, 44:418–436. doi: 10.1111/zsc.12108.

Ratnasingham S., Hebert P.D.N. 2007. BOLD: The Barcode of Life Data System (http://www.barcodinglife.org). Molecular Ecology Notes, 7:355–364. doi.

Rota J., Twort V.G., Chiocchio A., Peña C., Wheat C.W., Kaila L., Wahlberg N. 2022. The unresolved phylogenomic tree of butterflies and moths (Lepidoptera): potential causes and consequences. Syst. Entom., 47:531–550. doi: 10.1111/syen.12545.

Rowe K.C., Singhal S., Macmanes M.D., Ayroles J.F., Morelli T.L., Rubidge E.M., Bi K., Moritz C.C. 2011. Museum genomics: low-cost and high-accuracy genetic data from historical specimens. Mol. Ecol. Res., 11:1082–1092. doi.

Savva G., Dicks J., Roberts I.N. 2003. Current approaches to whole genome phylogenetic analysis. Briefings in Bioinformatics, 4:63–74. doi: 10.1093/bib/4.1.63.

Schmieder R., Edwards R. 2011. Quality control and preprocessing of metagenomic datasets. Bioinformatics. Bioinformatics, 27:863–864. doi: 10.1093/bioinformatics/btr026.

Sproul J.S., Maddision D.R. 2017. Sequencing historical specimens: successful preparation of small specimens with low amounts of degraded DNA. Mol. Ecol. Res., 17:1183–1201. doi: 10.1111/1755-0998.12660.

St Laurent R.A., Hamilton C.A., Kawahara A.Y. 2018. Museum specimens provide phylogenomic data to resolve relationships of sack-bearer moths (Lepidoptera, Mimallonoidea, Mimallonidae). Syst. Entom., 43:729–761. doi: 10.1111/syen.12301.

Staats M., Erkens R.H.J., Vossenberg B.v.d., Wieringa J.J., Kraaijeveld K., Stielow B., Geml J., Richardson J.E., Bakker F.T. 2013. Genomic treasure troves: complete genome sequencing of herbarium and insect museum specimens. PLoS One, 8:e69189. doi.

Suyama M., Torrents D., Bork P. 2006. PAL2NAL: robust conversion of protein sequence alignments into the corresponding codon alignments. Nucleic Acids Research, 34:W609–W612. doi: 10.1093/nar/gkl315.

Toussaint E.F.A., Breinholt E.W., Earl C., Warren A.D., Brower A.V.Z., Yago M., Dexter K.M., Espeland M., Pierce N.E., Lohman D.J., et al. 2018. Anchored phylogenomics illuminates the skipper butterfly tree of life. BMC Evolutionary Biology, 18:101. doi: 10.1186/s12862-018-1216-z.

Twort V.G., Minet J., Wheat C.W., Wahlberg N. 2021. Museomics of a rare taxon: placing Whalleyanidae in the Lepidoptera Tree of Life. Syst. Entom., 46:926–937. doi: 10.1111/syen.12503.

Wahlberg N., Wheat C.W. 2008. Genomic outposts serve the phylogenomic pioneers: designing novel nuclear markers for genomic DNA extractions of Lepidoptera. Syst. Biol., 57:231–242. doi: 10.1080/10635150802033006.

Wang H., Holloway J.D., Wahlberg N., Wang M., Nylin S. 2019. Molecular phylogenetic and morphological studies on the systematic position of Heracula discivitta reveal a new subfamily of Pseudobistonidae (Lepidoptera: Geometroidea). Syst. Entom., 44:211–225. doi.

Wang Z., Gerstein M., Snyder M. 2009. RNA-Seq: a revolutionary tool for transcriptomics. Nature Reviews Genetics, 10:57–63. doi.

Wei C.-H., Yen S.-H. 2017. Mimaporia, a new genus of Epicopeiidae (Lepidoptera), with description of a new species from Vietnam. Zootaxa, 4254:537–550. doi: 10.11646/zootaxa.4254.5.3.

Willerslev E., Cooper A. 2005. Ancient DNA. Proc. R. Soc. B, 272:3–16. doi: 10.1098/rspb.2004.2813.

Zerbino D.R., Birney E. 2008. Velvet: Algorithms for de novo short read assembly using de Bruijn graphs. Genome Res., 18:821–829. doi.

Zhang J., Cong Q., Shen J., Brockmann E., Grishin N.V. 2019. Genomes reveal drastic and recurrent phenotypic divergence in firetip skipper butterflies (Hesperiidae: Pyrrhopyginae). Proc. R. Soc. B, 286:20190609. doi.

Zhang Y.Y., Huang S.Y., Liang D., Wang H.S., Zhang P. 2020. A multilocus analysis of Epicopeiidae (Lepidoptera, Geometroidea) provides new insights into their relationships and the evolutionary history of mimicry. Molecular Phylogenetics and Evolution, 149:106847. doi: 10.1016/j.ympev.2020.106847.

