## Supplementary figures and images for "One Method to Sequence Them All? Comparison between Whole-Genome Sequencing (WGS) and Target Enrichment (TE) of museum specimens from the moth families Epicopeiidae and Sematuridae (Lepidoptera)"

### FigureS1

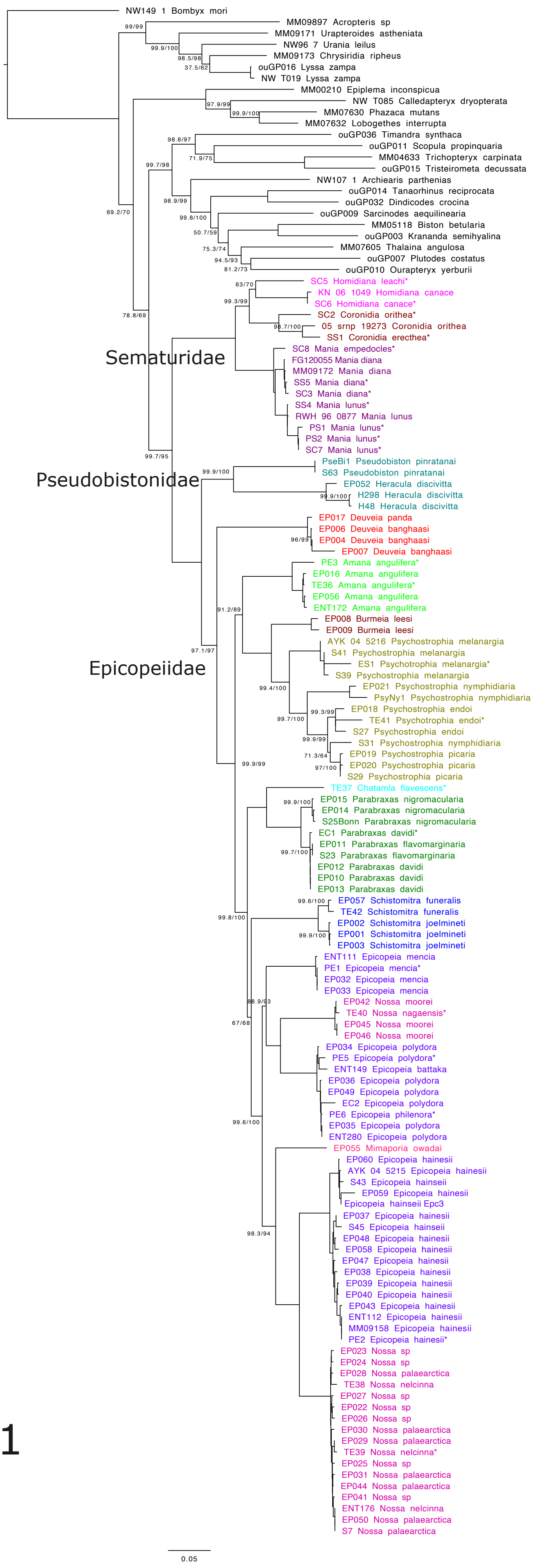
